# Monocarboxylate Transporter 1 (MCT1) mediates succinate export in the retina

**DOI:** 10.1101/2021.11.19.469314

**Authors:** Celia M. Bisbach, Daniel T. Hass, James B. Hurley

## Abstract

**Purpose:** Succinate is exported by the retina and imported by eyecup tissue. The transporter(s) mediating this process have not yet been identified. Recent studies showed that Monocarboxylate Transporter 1 (MCT1) can transport succinate across plasma membranes in cardiac and skeletal muscle. Retina and retinal pigment epithelium (RPE) both express multiple MCT isoforms including MCT1. We tested the hypothesis that MCTs facilitate retinal succinate export and RPE succinate import.

**Methods:** We assessed retinal succinate export and eyecup succinate import in short term *ex vivo* culture using gas chromatography-mass spectrometry. We test the dependence of succinate export and import on pH, proton ionophores, conventional MCT substrates, and the MCT inhibitors AZD3965, AR-C155858, and diclofenac.

**Results:** Succinate exits retinal tissue through MCT1 but does not enter RPE through MCT1 or any other MCT. Intracellular succinate levels are a contributing factor that determines if an MCT1-expressing tissue will export succinate.

**Conclusions:** MCT1 facilitates export of succinate from retinas. An unidentified, non-MCT transporter facilitates import of succinate into RPE.

## Introduction

Succinate has a unique role in the eye. It is a valuable metabolic fuel, and evidence supports a model where succinate produced by fumarate respiration and exported from the retina is oxidized by the neighboring retinal pigment epithelium (RPE).^1,2^ Succinate is also implicated in retinal pathology. In a model of O_2_-induced retinopathy, succinate accumulates in the retina and drives angiogenesis via SUCNR1 signaling on retinal ganglion cells.^3^ SUCNR1-deficiency in mice promotes premature sub-retinal dystrophy, and SUCNR1 variants are associated with age-related macular degeneration in a human population.^4^ Under both normal and pathological circumstances, succinate mediates its effects in a non-cell autonomous manner. However, the transporter used by succinate to exit or enter any ocular cell type has not yet been identified.

Monocarboxylate carrier 1 (MCT1) can facilitate succinate export in exercising muscle and ischemic heart.^5,6^ Retinas express both MCT1 and MCT4, while RPE cells in eyecups express MCT1 and MCT3.^7,8^ Based off this, we set out to test if MCT1 might also be responsible for facilitating retinal succinate export and/or RPE succinate import, and whether additional MCTs also transport succinate in the eye.

## Results

### MCT1 mediates succinate export in retinas

Succinate is an abundant metabolite in retinas. Retinas in short-term *ex vivo* culture export a molar amount of succinate equivalent to their entire intracellular succinate pool approximately every 40 minutes (**Figure 1A** and **Figure 1B**).^1^ In order to evaluate the role of MCTs in retinal succinate export, MCT activity must be reduced. Retinas are exceptionally glycolytic, so reducing MCT activity could block flux through glycolysis and indirectly affect intracellular succinate production rather than its release. To control for this potentially confounding factor, we supplied retinas with 5 mM ^12^C-glucose and 50 μM ^13^C-malate in all experiments testing the role of MCTs in succinate export in this section (**Figure 1C**). Including this tracer quantity of ^13^C-malate allows for quantification of succinate produced both by oxidative (m0 succinate) and by reductive (m4 succinate) TCA cycle activity.^1^ m4 succinate production does not depend on acetyl-CoA availability, so by monitoring rates of both m0 and m4 succinate export, we can attribute a decrease in succinate export to specific inhibition of MCT-mediated succinate transport, not just a change in intracellular succinate production.

**Figure 1:**
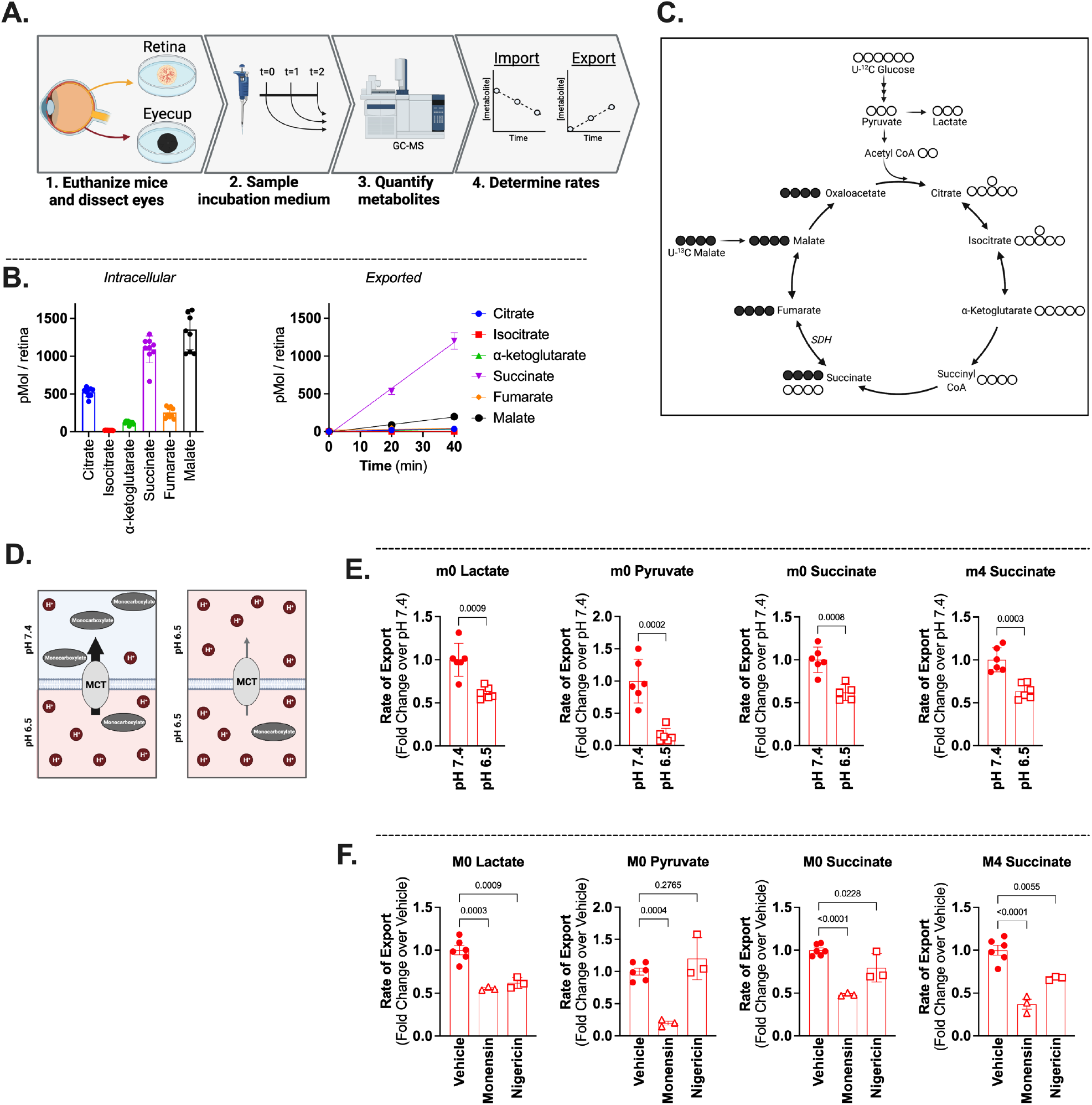
H^+^ dependence of succinate export in retinas. **(A)** Experimental workflow for determination of metabolite export and import rates. **(B)** Diagram showing the isotopologues of succinate which are made by oxidative metabolism of U-^12^C-glucose and U-^13^C-succinate. Open circles represent ^12^C, filled circles represent ^13^C. **(C)** Intracellular levels of TCA cycle metabolites from freshly dissected retinas (left, n = 9) and rate at which they are exported when retinas are incubated in 5 mM glucose(right, n = 6). **(D)** Representation of how pH gradients can influence MCT activity. **(E)** Rate of export of the canonical MCT substrate m0 lactate and m0 pyruvate, as well as m0 succinate and m4 succinate from retinas incubatd in 5 mM ^12^C-glucose and 50 μM ^13^C-malate. p-values determined using Welch’s t test (n = 6 retinas). **(F)** Rate of export of m0 lactate, m0 pyruvate, m0 succinate, and m4 succinate from retinas in the presence of 100 μM monensin or 100 μM nigericin. p-values determined using a one-way ANOVA (n = (6, vehicle) (3, monensin) (3, nigericin)).

MCT-dependent substrate transport depends on pH because it transports protons along with each anionic substrate. Lowering pH on the *trans* (in this case extracellular) side of a MCT decreases its ability to export anions, as it must now transport protons against a concentration gradient (**Figure 1D**). We measured succinate export in retinas incubated in medium adjusted to pH 7.4 or 6.5 and observed that lower extracellular pH reduced export of m0 and m4 succinate, as well as the canonical MCT substrates lactate and pyruvate (**Figure 1E)**. We treated retinas incubated in pH 7.4 medium with H^+^ ionophores (the H^+^/Na^+^ ionophore monensin or the H^+^/K^+^ ionophore nigericin) to diminish any proton gradient that may normally drive MCT-mediated export in retinas. Rates of m0 lactate, m0 pyruvate, m0 succinate, and m4 succinate export all were diminished by the ionophores (**Figure 1F**).

We next tested several MCT isoform-specific inhibitors to determine if we could identify which specific MCT(s) facilitate succinate export in retinas (**Table 1**). The dual MCT1/MCT2 inhibitors AR-C155858 or AZD3965 (**Figure 2A** and **Figure 2B**) slow m0 and m4 succinate export in a dose-dependent manner. These effects are likely mediated through inhibition of MCT1, since MCT2 appears either to not be expressed or to be expressed only at very low levels in the retina.^9–11^ The dual MCT1/MCT4 inhibitor diclofenac (**Figure 2C**) also slows both m0 and m4 succinate export. It is possible that the increased effectiveness of diclofenac relative to the dual MCT1/MCT2 inhibitors is due to inhibition of MCT4 in addition to MCT1. However, in cultured myotubules expressing MCTs 1, 2, and 4, reducing MCT4 expression did not alter succinate export.^5^ Since diclofenac is structurally distinct from AZD3965 and AR-C155858, its increased effectiveness could also be due to an altered mechanism of action or increased permeability. As a result, these experiments show that MCT1 is responsible for a portion of succinate export in retinas and MCT4 may transport a portion of succinate in retinas.

**Table 1:** Reported kinetic parameters for various MCT inhibitors. “n.d.” indicates no data for the effectiveness of an inhibitor for that MCT has been reported. *no inhibition observed up to 10 μM †no inhibition observed up to 1 μM, some inhibition observed at 10 μM ‡ The K_i_ for diclofenac in a mixed population of MCT1, MCT2, and MCT4 is 20 μM

| Inhibitor | *MCT1* | *MCT2* | *MCT3* | *MCT4* | references |
| --- | --- | --- | --- | --- | --- |
| AZD3965 | 3.2 nM ($K_i$) | 20 nM ($K_i$) | No Inhibition | No inhibition* | 16 |
| AR-C155858 | 2.3 nM ($K_i$) | 10 nM | n.d. | Minimal inhibition † | 18 |
| Diclofenac‡ | 1.45 μM ($IC_{50}$) | n.d. | n.d. | 0.14 μM ($IC_{50}$) | 19,32 |

**Figure 2:**
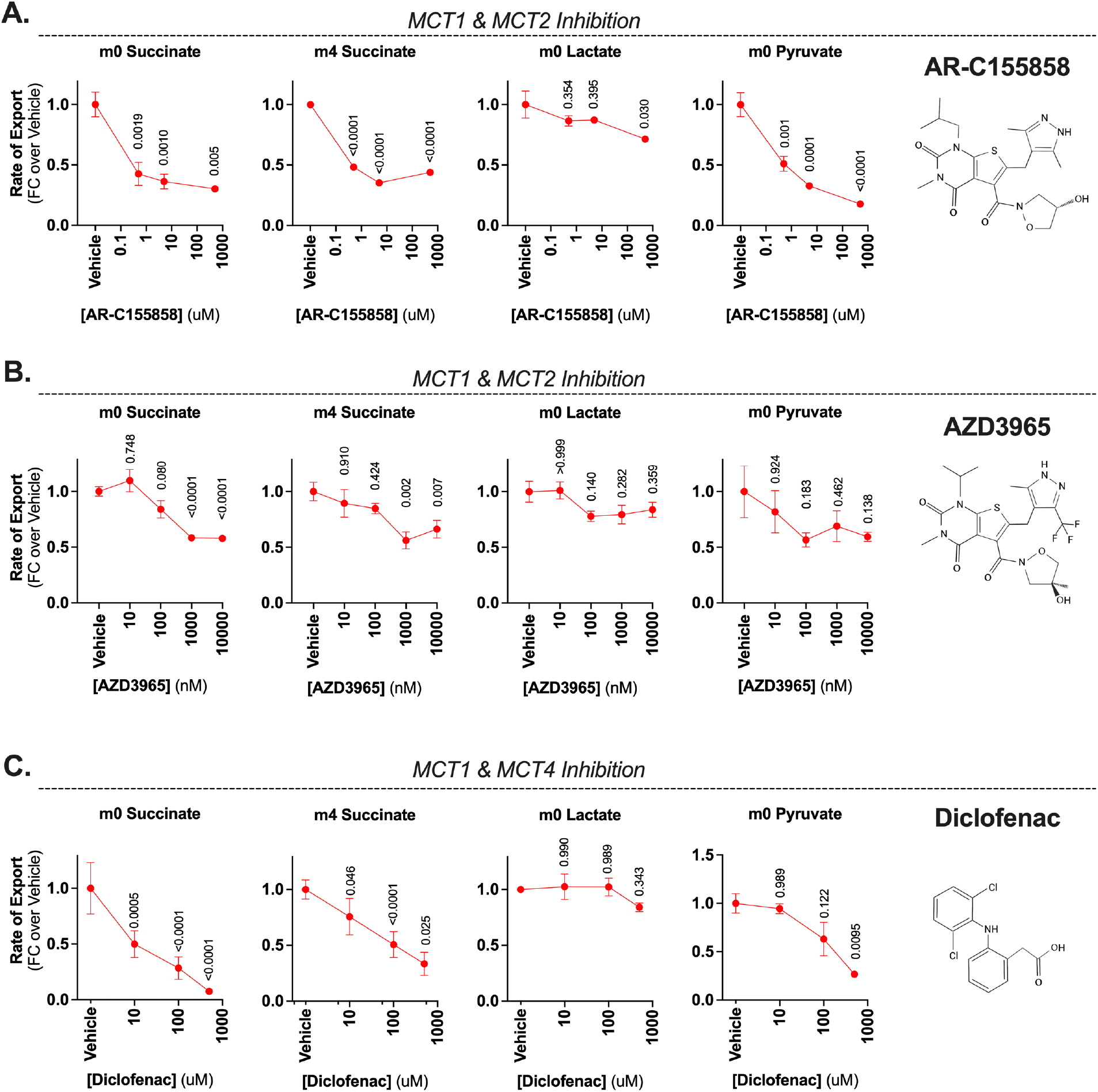
Effect of MCT inhibition on succinate export in retinas. **(A**,**B**,**C)** Influences of various MCT inhibitors on retinal export of m0 succinate, m4 succinate, m0 lactate and m0 pyruvate. Rates determined by sampling incubation media from retinas incubated in 5 mM ^12^C-glucose and 50 μM ^13^C-malate. P-values shown above each point were calculated using an ordinary one-way ANOVA (all conditions compared to vehicle) followed by Dunnett’s correction for multiple comparisons. **(A)** AR-C155858 (n = (3, 0 nM) (3, 500 nM) (3, 5 μM) (3, 500 μM). **(B)** AZD3965 (n = (9, 0 nM) (3, 10 nM) (9, 100 nM) (6, 1 μM) (9, 10 μM)). **(C)** Diclofenac (n = (6, 0 nM) (3, 10 uM) (6, 100 uM) (3, 500 μM).

### Eyecup succinate import is not MCT mediated

Unlike retina tissue, eyecup explants do not export succinate and instead measurably deplete it from the incubation (**Figure 3A**).^1,2^ Eyecup tissue is composed of sclera, choroidal endothelial cells, and RPE cells. MCT expression on RPE cells is polarized, with MCT3 localized to the basal RPE surface and robust MCT1 expression on the apical processes.^8,10,12^ Because apical processes are in direct contact with the succinate-exporting retina, we hypothesized that MCT1 may be responsible for both retina succinate export and RPE succinate import.

**Figure 3:**
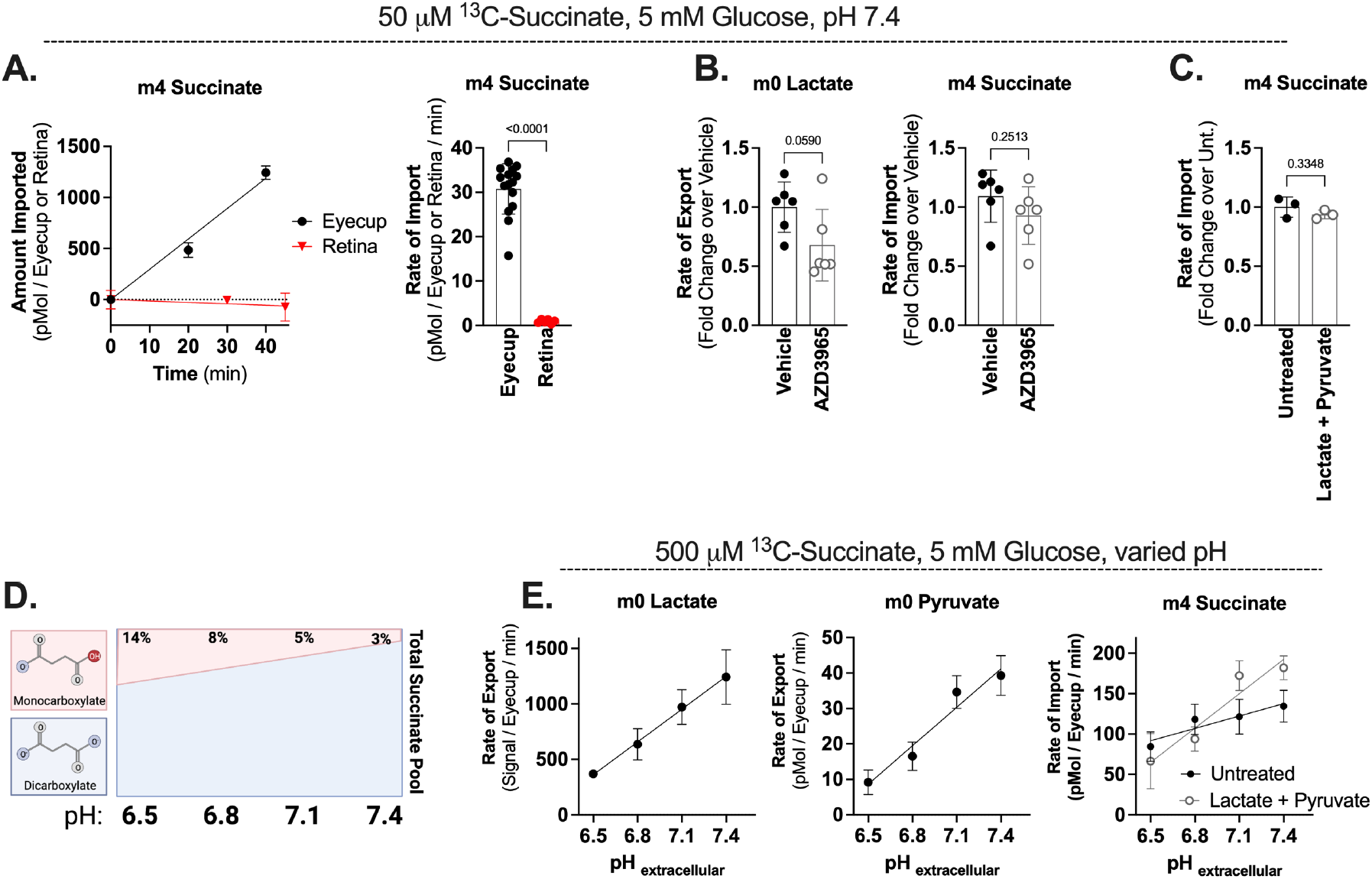
MCTs do not import succinate in eyecups. **(A)** Depletion of succinate from the incubation media by eyecups (n=14) and retinas (n=6) incubated in 5 mM ^12^C-glucose and 50 μM ^13^C-succinate. Linearity of import over time is shown on the left, calculated rates of import shown on the right. **(B)** Rates of lactate export and succinate import in the presence of 100 μM AZD3965. p-value determined by a two-tailed Welch’s t-test (n=6 eyecups for each condition). **(C)** Rate of succinate import in the presence of 15 mM Lactate and 1 mM Pyruvate. P-value determined by a two-tailed Welch’s t-test (n=3 eyecups for each condition). Exogenous lactate and pyruvate prevent determination of the rates of lactate or pyruvate export in this experiment. **(D)** Graphical representation of how pH can influence the fraction of monocarboxylate succinate. The ratio of of monocarboxylate:dicarboxylate succinate was determined using the Henderson-Hasselbalch equation and a pKa of 5.69. **(E)** Rate of succinate import by eyecups supplied with 5 mM ^12^C-glucose and 500 μM ^13^C-succinate, in the presence or absence of 15 mM lactate and 1 mM pyruvate. (n=6 untreated eyecups at each pH, 3 lactate + pyruvate eyecups at each pH).

We first tested if MCT1 is responsible for eyecup succinate import by measuring import of 50 μM U-^13^C-succinate from pH 7.4 incubation medium in the presence of the MCT1 inhibitor AZD3965 (**Figure 3B**). AZD3965 suppressed lactate export but did not inhibit succinate import. However, it is possible that MCT3 maintains succinate uptake when MCT1 is inhibited. We were unable to identify a well-characterized MCT3 inhibitor, so we tested if we could outcompete U-^13^C-succinate import with an excess of lactate and pyruvate (substrates for both MCT1 and MCT3 which would act as competitive inhibitors) (**Figure 3C**). Lactate and pyruvate did not alter the rate of succinate import.

We next considered that eyecups could express a high-affinity, non-MCT succinate transporter, and that MCTs might only become engaged in succinate import under specific conditions (such as when pH is low and/or when extracellular succinate concentrations are higher). Lowering pH in *cis* with succinate could modulate MCT-mediated succinate transport in two ways: (1) as shown previously in **Figure 1D**, a high proton gradient can drive transport, and (2) at acidic pH, a greater fraction of succinate exists as a MCT-transportable monocarboxylate (**Figure 3D**).^5^ With this in mind, we tested if we could engage measurable MCT-mediated succinate import in eyecups by acidifying the extracellular pH and increasing the extracellular succinate concentration (**Figure 3E**). As expected, the rates of lactate and pyruvate export decreased as extracellular pH was acidified and MCTs began to export these substrates against an increasingly steep proton gradient. However, the rate of succinate import also decreased as pH acidified, indicating that the steeper proton gradient was ineffective at enhancing succinate import.

It is possible that low pH suppresses mitochondrial succinate oxidation, and that an increase in MCT-mediated succinate transport is masked by a decrease in succinate oxidation at acidic pH. To control for this, we tested if the MCT substrates lactate and pyruvate might be able to outcompete a greater fraction of succinate uptake as pH acidified (**Figure 3E**). Even at the lowest pH, lactate and pyruvate had no significant influence on succinate import. This indicates that an unidentified succinate transporter and not MCT1 or MCT3 is the primary succinate importer in eyecups.

### Eyecups can be induced to export succinate via MCT1

Although both retina and eyecup express MCT1, retinal explants export succinate while eyecup explants do not. In fact, MCT1 can be found on red skeletal muscle, cardiac muscle, red blood cells, liver, kidney cortex and tubule cells, adipose tissue, cerebral neurons and glia, and pancreatic acinar cells, but retina and pancreas are the only tissues reported to date that export succinate under basal conditions ^8,9,13,14^. We next sought to determine if we could identify a metabolic difference between MCT1-expressing tissues in the eye which causes succinate export to become engaged.

In other tissues that can be induced to export succinate (such as exercising muscle or ischemic heart tissue), succinate export is accompanied by a rise in intracellular succinate levels and intracellular acidification.^5,6,15^ We compared intracellular succinate levels between retina and eyecup tissue and saw that retinas contain approximately 20-fold more succinate per μg protein compared to eyecups (**Figure 4A**).

**Figure 4:**
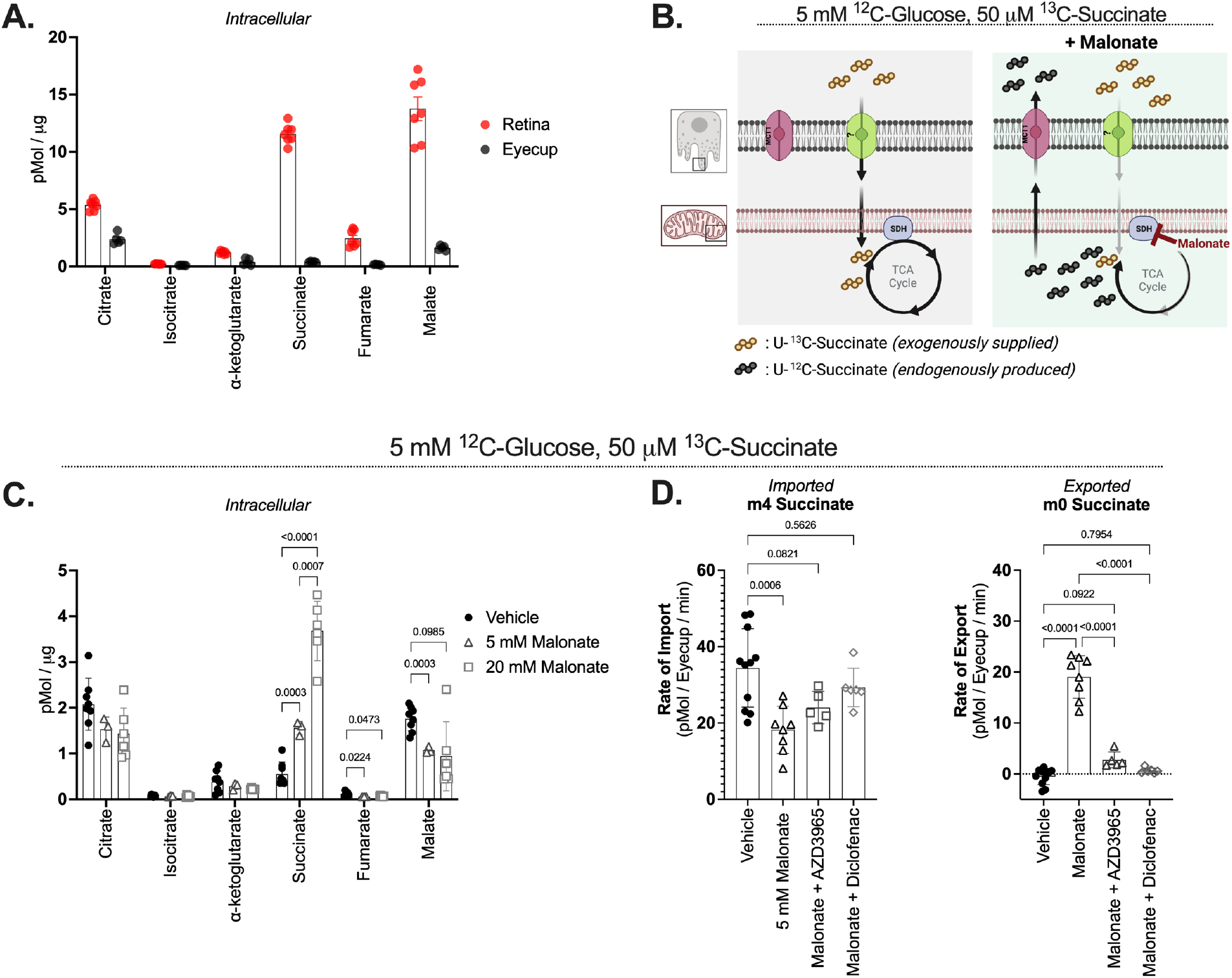
Eyecups can be induced to export succinate via MCT1. **(A)** Total intracellular metabolites per μg dry protein in freshly dissected retinas and eyecups (n=7 retinas and 5 eyecups). **(B)** Influences of malonate on succinate dynamics in eyecups. **(C)** Intracellular metabolite levels per μg dry tissue in eyecups incubated in 0, 5, or 20 mM malonate for 40 minutes. P-values determined using an ordinary one-way ANOVA followed by Dunnett’s correction for multiple comparisons. (n = (8, Vehicle) (3, 5 mM) (6, 20 mM)). **(D)** Rates of m4 succinate import and m0 succinate export from eyecups incubated with 5 mM malonate, 5 mM malonate + 100 nM AZD3965, or 5 mM malonate + 100 μM diclofenac. P-values determined using a one-way ANOVA followed by Dunnett’s correction for multiple comparisons. (n = (11, vehicle) (8, malonate) (5, malonate + AZD3965) (6, malonate + diclofenac)).

To test if increasing eyecup succinate levels were sufficient to induce succinate export, we treated eyecups with the SDH inhibitor malonate and observed a dose-dependent increase in intracellular succinate levels (**Figure 4B** and **4C**). When we sampled incubation media from retinas incubated with malonate, we observed that eyecups stopped importing exogenous succinate (m4) and began to export endogenous succinate (m0) (**Figure 4D**). This succinate export was effectively shut down by the MCT1 inhibitors AZD3965 and diclofenac (**Figure 4D**). The nearly complete inhibition of malonate-induced m0 succinate export by AZD3965 in eyecups indicates that MCT3 is unable to transport succinate, since AZD3965 has no inhibitory effect on MCT3 even well above the concentration we used.^16^ Taken together, these results show that a high intracellular succinate concentration is at least one contributing factor that contributes to a tissue exporting succinate through MCT1 **(Figure 4B**).

## Discussion

The goal of this study was to understand how succinate is transported between cells in an eye. Our results indicate that MCT1 can be added to the model of succinate transit in the retinal ecosystem as the retinal succinate exporter, but not as the RPE succinate importer.

MCT inhibitors are ineffective at reducing lactate export in retinas. The MCT1/2 inhibitor AZD3965 is a competitive inhibitor, so its increased effectiveness at reducing pyruvate export relative to lactate export could be due in part to the exceptionally high intracellular lactate levels in photoreceptors.^17^ The MCT1/2 inhibitor AR-C155858 and the MCT1/4 inhibitor diclofenac both appear to be non-competitive inhibitors and so their effectiveness should not depend on lactate concentration.^18,19^ Consistent with this, AR-C155858 exhibited the best dose-dependent inhibition of lactate efflux, although pyruvate efflux was still more strongly inhibited. Non-ionic diffusion of the free acid form of lactate across the plasma membrane could be responsible for a portion of the sustained lactate export in the presence of MCT inhibitors. The free diffusion of lactate is reported to be faster than that of pyruvate, and retinas have the high intracellular lactate concentrations required to drive this process.^20–22^ Retinas may also express an inhibitor-insensitive transporter capable of facilitating lactate export in the absence of MCT function.

Determining why MCT1 plays no significant role in importing succinate under physiological *ex vivo* conditions in both retinas and eyecups is important to understanding the biological implications of circulating succinate. The current understanding of MCT activity states that at equilibrium, there will be a balance between the [H^+^]_out,_ [monocarboxylate]_out_ and [H^+^]_in,_ [monocarboxylate]_in_.^23^ A recent report has illustrated the added influence pH has on modulating the amount of MCT-transportable succinate due to its physiologically relevant pKa (**Figure 3D**).^5^ Based off this, we reasoned that we might detect MCT1-mediated succinate import in eyecups if we enhanced it by lowering the extracellular pH and supplying a higher concentration of extracellular succinate (**Figure 3E**). However, this did not enhance succinate import. Perhaps an unidentified succinate importer has a significantly higher affinity for succinate than MCT1 does, and import of succinate into eyecups may not be the rate limiting step for overall succinate oxidation. In this scenario, any change to the small amount of succinate that could be imported by MCT1 would not affect the overall process of succinate depletion and thus we would not be able to detect it in our assay.

Even though retinas express copious MCT1, they appear to be incapable of using it as a succinate importer. Retinas incubated in U-^13^C-succinate do not deplete measurable amounts from the media (**Figure 3A**) and intracellular downstream intermediates are ^13^C-labeled by succinate only to a minor degree.^1,24^ Since a small fraction of downstream intermediates can be labeled by exogenous succinate in retinas, MCT1 may be able to import a small amount of succinate. However, since supplying retinas with as much as 100 mM succinate does not stimulate significant O_2_ consumption, it is unlikely that the small amount of MCT1-mediated succinate import in retinas is biologically relevant.^2^

Considering that for a MCT at equilibrium [H^+^]_out,_ [monocarboxylate]_out_ = [H^+^]_in,_ [monocarboxylate]_in._, the likely explanation for the observed unidirectional MCT1 activity in retinas is that they have both high intracellular succinate levels (**Figure 4A**) and an unusually acidified cytosol. *In vivo* pH measurements made from a cat eye report the pH of the retina to be ∼7.1, while the pH of the choroid (near the RPE) was ∼7.4.^25^ A retina maintains an unusually high rate of aerobic glycolysis relative to the RPE (and most other tissues), which likely contributes to retinal acidification.^26,27^ The retina also produces succinate via both oxidative TCA cycle activity and reduction of fumarate, which could contribute to maintaining high intracellular succinate levels.^1^ This may explain why most MCT1-expressing tissues do not export succinate under basal conditions. Since most tissues do not rely on aerobic glycolysis and fumarate respiration to the same extent that the retina does, they lack the requisite acidified cytosol and high intracellular succinate levels required to engage MCT1-mediated succinate transport.

Like healthy retina tissue, MCT1 overexpressing cancers also rely on aerobic glycolysis and convert a large fraction of the glucose they consume to lactate.^28^ Succinate has recently been identified as a metabolite exported by cancer cells and that promotes metastatic activity through SUCNR1 signaling.^29^ Determining if this succinate also is exported is via MCT1 will allow it to be targeted for inhibition by existing MCT1-specific chemotherapy drugs.

By identifying the retinal succinate exporter, we hoped we would be able to harness it to study the role of succinate exchange in the retinal ecosystem by enhancing or decreasing its expression. Unfortunately, the canonical role of MCT1 as a lactate and pyruvate transporter is fundamental to retinal health and it would be difficult to parse any succinate-specific phenotype from disruptions caused by inhibition of lactate and pyruvate transport. However, this general strategy may still be pursued once the RPE succinate importer has been identified.

## Methods

### Animals

All experiments used 2-6 month-old male and female wild-type C57BL6/J mice, either obtained from Jackson Labs or bred in house. These mice were housed at the SLU 3.1 facility at an ambient temperature of 25°C, with a 12-hour light cycle and ad libitum access to water and normal rodent chow. Experiments using animals conform to the ARVO guidelines for the use of animals in ophthalmic research.

### *Ex vivo* metabolite uptake/export

In all *ex vivo* metabolic analysis experiments, mice were euthanized by awake cervical dislocation and retinas and/or eyecups were dissected in Hank’s Buffered Salt Solution (HBSS; GIBCO, Cat#: 14025-076). Tissue was incubated in Krebs-Ringer buffer (formulations used in each figure specified below below) supplemented with 5 mM glucose and [U-^13^C]-succinic acid (Cambridge isotope CLM-1571-0.1) or [U-^13^C]-malate (Cambridge isotope CLM-8065) as indicated in each figure. For experiments using KRB buffer, buffer was pre-equilibrated at 37°C, 21% O_2_, and 5% CO_2_ prior to incubations and incubations were carried out at those conditions. For experiments where pH was modulated KRM buffer was used, buffer was pre-equilibrated at 37°C and room oxygen and incubations were carried out under those conditions. For determination of metabolite import or export rates, incubation media was sampled at 3 timepoints (typically 0, 20, and 40 minutes) and export or import was confirmed to be linear over time. Retinas were incubated in 200 μL and eyecups in 100 μL over this range of time. Inhibitors used were AZD3965 (Cayman Chemical no. 19912), AR-C155858 (MedChemExpress HY-13248), diclofenac sodium salt (Cayman Chemical no. 70680). Ethanol was used as a solvent for AZD3965, DMSO was used as a solvent for AR-C155858, and separate experiments were done using both DMSO and ethanol as a solvent for diclofenac.

### Buffer formulations used

Krebs-Ringer Bicarbonate (KRB) buffer was used in all experiments except for Figures 1E and 3E: (98.5 mM NaCl, 5.1 mM KCl, 1.2 mM KH_2_PO_4_, 1.2 mM MgSO_4_-7H_2_O, 2.7 mM CaCl_2_-2H_2_O, 20.8 mM HEPES, and 25.9 mM NaHCO_3_). Krebs-Ringer MOPS (KRM) buffer was used in Figures 1E and 3E: (98.5 mM NaCl, 5.1 mM KCl, 1.2 mM KH_2_PO_4_, 1.2 mM MgSO_4_-7H_2_O, 2.7 mM CaCl_2_-2H_2_O, 10 mM HEPES, and 15 mM MOPS). KRB buffer equilibrates to pH 7.4 at 37°C in a 5% CO_2_ incubator. KRM buffer was adjusted to the desired pH using HCl at 37°C in room O_2_.

### Metabolite Extraction

Media samples were added directly to 90% MeOH supplemented with 10 μM methylsuccinate and immediately lyophilized. Dried samples were stored at −80°C until derivatization. Metabolites were extracted from retina or eyecup tissue using 150 μL ice-cold 80% MeOH, 20% H_2_O supplemented with 10 μM methylsuccinate (Sigma, M81209) as an internal standard to adjust for any metabolite loss during the extraction and derivatization procedures. Tissues were disrupted by sonication and incubated on dry ice for 45 minutes to precipitate protein. Protein and cell debris was pelleted at 17,000 x g for 30 minutes at 4°C. The supernatant containing metabolites was lyophilized at room-temperature until dry and stored at −80°C until derivatization. The protein pellet was resuspended by sonication in RIPA buffer (150 mM NaCl, 1.0% Triton X-100, 0.5% sodium deoxycholate, 0.1% SDS, 50 mM Tris, pH 8.0) and the amount of protein was determined by a BCA assay (ThermoFisher, 23225).

### Metabolite Derivatization

Lyophilized samples were first derivatized in 10 μL of 20 mg/mL methoxyamine HCl (Sigma, Cat#: 226904) dissolved in pyridine (Sigma, Cat#: 270970) at 37°C for 90 minutes, and subsequently with 10 μL tert-butyldimethylsilyl-N-methyltrifluoroacetamide (Sigma, Cat#: 394882) at 70°C for 90 minutes.

### Gas Chromatography-Mass Spectrometry

Metabolites were analyzed on an Agilent 7890/5975C GC-MS using selected-ion monitoring methods described in previous work.^30^ Peaks were integrated in MSD ChemStation (Agilent), and correction for natural isotope abundance was performed using IsoCor software.^31^ Corrected metabolite signals were converted to molar amounts by comparing metabolite peak abundances in samples with those in a standard mix (which was prepared to contain known quantities of metabolites and run along each individual experiment). These known metabolite concentrations were used to generate a standard curve which allowed for metabolite quantification.

### Statistical Analysis

Statistical tests were performed in GraphPad Prism v9. The test used is indicated in each figure. Briefly, Welch’s t-test was used in instances where two groups were compared, and an ordinary one-way ANOVA followed by the recommended correction for multiple comparison was used when multiple conditions where compared. “n” is indicated in each figure legend and represents biological replicates (i.e. n = 3 indicates that 3 rate measurements were obtained from 3 different retinas, rather than 3 rate measurements made from the same retina). Retinas and eyecups from the same mouse were always used in different experimental conditions and thus were considered biological replicates.

## Acknowledgements

The authors who contributed to this work are CMB, DTH, and JBH. CMB conceptualized the work with helpful discussions from JBH and DTH. CMB performed the experiments, analyzed the data, and prepared the manuscript. JBH and DTH assisted with editing the manuscript. We thank Martin Sadilek (UW Chemistry) for assistance with maintaining GC-MS instrumentation and methods, and Whitney Cleghorn and Marcos Nazario for mouse colony maintenance. Funding was provided by F31EY031165 (CMB), 5T32EY007031-42 (DTH), and EY06641 and EY017863 (JBH).

